# Gut microbiota-derived tryptamine impairs insulin sensitivity

**DOI:** 10.1101/2022.03.05.483098

**Authors:** Lixiang Zhai, Haitao Xiao, Chengyuan Lin, Yan Y. Lam, Hoi Leong Xavier Wong, Mengxue Gong, Guojun Wu, Yusheng Deng, Ziwan Ning, Chunhua Huang, Yijing Zhang, Min Zhuang, Chao Yang, Eric Lu Zhang, Ling Zhao, Chenhong Zhang, Xiaodong Fang, Wei Jia, Liping Zhao, Zhao-xiang Bian

## Abstract

Gut-microbiota plays a pivotal role in development of type 2 diabetes (T2D), yet the molecular mechanism remains elusive. Here, we show that tryptamine, a microbial metabolite of tryptophan, impairs glucose tolerance and insulin sensitivity. Tryptamine presents a higher level in monkeys with spontaneous diabetes and human with T2D and positively correlated with the glucose tolerance. In parallel, tryptamine level was suppressed by dietary fibers intervention in T2D subjects and negatively correlated with improvement of glucose tolerance. The inhibitory effect of tryptamine on insulin signaling as shown was dependent on a trace amine-associated receptor 1 (TAAR1)-extracellular signal-regulated kinase (ERK) signaling axis. Monoassociation of T2D-associated tryptamine-producing bacteria *Ruminococcus gnavus* impairs insulin sensitivity in pseudo germ-free mice. Our findings indicate gut microbiota-derived tryptamine contributes to the development of insulin resistance in T2D and may serve as a new target for intervention.

**Graphical Abstract:** 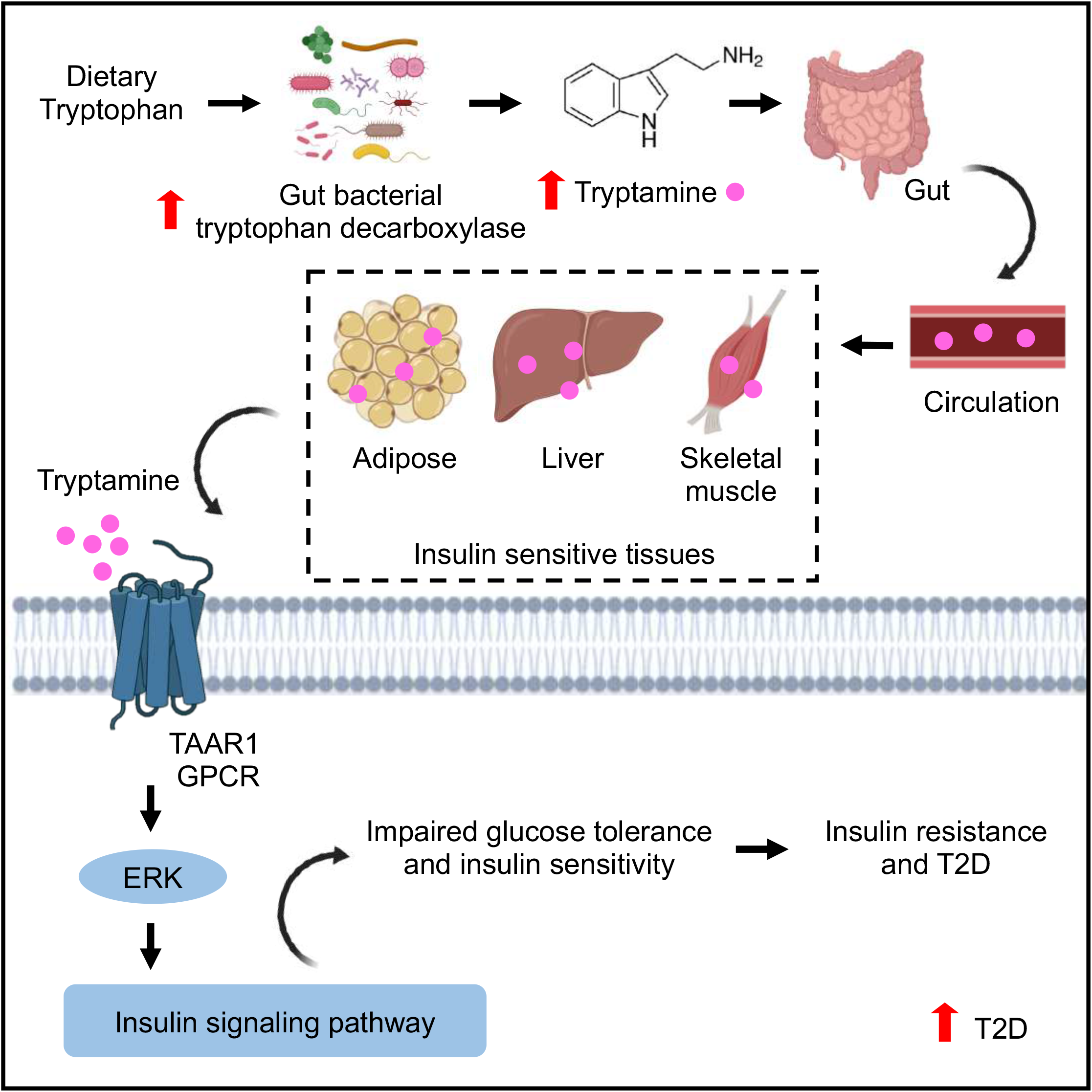

## Introduction

Type 2 diabetes (T2D) is a metabolic disorder characterized by high blood glucose levels due to insulin resistance and impaired insulin secretion (Galicia-Garcia et al., 2020). With the economic development and urbanization, T2D has become a major healthcare problem in both developing and developed countries. Over-consumption of unhealthy diets and sedentary lifestyles are the major factors that result in the development of T2D (Czech, 2017).

Gut microbiome has been extensively studied in in the past decades for its involvement in the pathophysiology of T2D (Gurung et al., 2020). Gut microbiota is capable of metabolizing nutrients and undigested food, thereby yielding microbial metabolites from carbohydrate fermentation, protein fermentation and metabolism of choline and cholesterol (Rooks and Garrett, 2016). Given that the composition of gut microbiota is dramatically changed in T2D, gut-microbial metabolites including lipopolysaccharides, short-chain fatty acids, bile acids, trimethylamine N-oxide and imidazole propionate are also significantly altered and exhibiting either beneficial or detrimental effects to glucose tolerance in both animal and human studies (Canfora et al., 2019; Prawitt et al., 2011).

Compared to the extensive research on these microbial metabolites, our current understandings of the pathophysiological relevance of microbial metabolites derived from protein fermentation on T2D remains limited. Dietary amino acids can be utilized and catabolized by gut microbiota into numerous metabolites that regulate important biological processes including intestinal barrier function, immune responses, and neurotransmission (Liu et al., 2020). Recent studies also identified aromatic and branched-chain amino acids derived microbial metabolites with bioactivities to affect intestinal permeability and systemic immunity (Dodd et al., 2017; Neis et al., 2015). However, there is limited knowledge into the role of amino acids-derived microbial metabolites in the pathogenesis of T2D. This study aimed to understand the role of gut bacteria and gut-microbial metabolites derived from amino acids in the development of T2D.

## Results

### Tryptamine is increased in monkeys with spontaneous diabetes and humans with type 2 diabetes

In the present study, we firstly employed crab-eating macaques (*Macaca fascicularis*) with spontaneous diabetes, a well-known pre-clinical model of T2D (Chen et al., 2018; Nagpal et al., 2018), to identify gut-microbial metabolites that may contribute to the development of T2D. Briefly, aged-matched male monkeys (n=78) were used based on their fasting blood glucose (FBG, mg/dL) and HbA1c (%) levels and were assigned to the normal group, pre-diabetes group and diabetes group (n=26 for each group) (Supplement Figure. 1A-D) according to the criteria references of diabetic studies on monkeys (Wang et al., 2019). We used fecal suspension from normal and diabetic monkeys to study the systemic effects of fecal suspension on glucose tolerance in HFD-fed mice. Notably, the HFD-mice fed with fecal suspension from diabetic groups exhibited a higher glucose level than the mice fed with fecal suspension from normal groups, indicating the fecal metabolites from diabetic monkeys impair glucose tolerance and aggravate the development of T2D (Figure.1A). Based on this finding, we conducted untargeted metabolomics to analyze the changes of fecal metabolome in monkeys from different groups (Supplement Figure.1E and Supplement Table.S1). Among all significantly changed fecal metabolites, we revealed tryptamine, a microbial metabolite from tryptophan metabolism, is positively correlated with the FBG and HbA1c index, suggesting tryptophan metabolism by gut microbiota is associated with spontaneous diabetes in monkeys and may be involved in the development of T2D (Figure. 1B and Supplement Table. S2).

**Figure.1.**
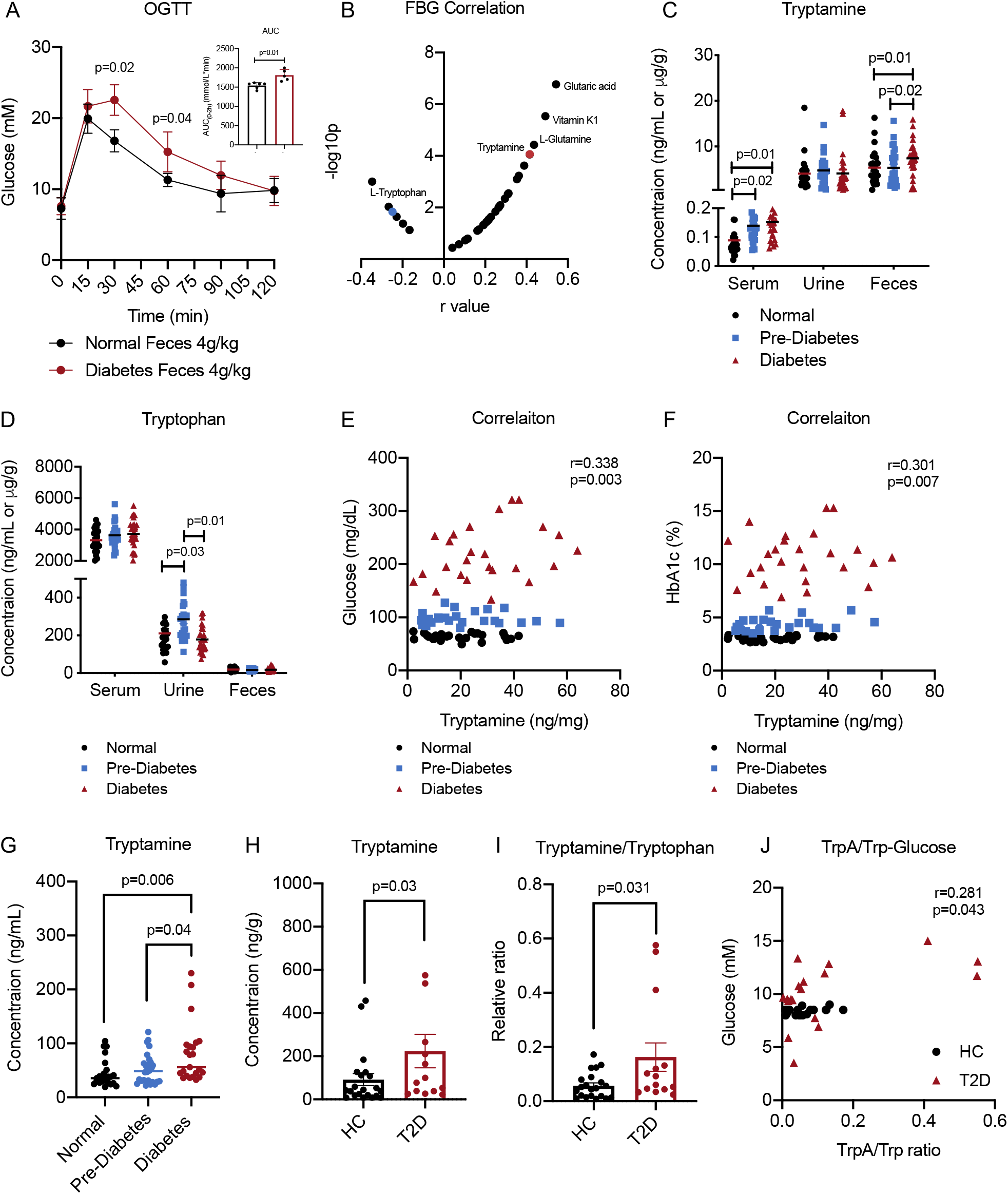
Tryptamine is increased in monkeys with spontaneous diabetes and human with type 2 diabetes. (A) Oral glucose tolerance test (OGTT) in HFD-fed mice after treatment of fecal suspension from normal group and diabetes group once per day for 5 days (n=6/group). (B) Spearman correlation of fecal metabolites and FBG level in diabetic monkeys. (C-D) Tryptamine and tryptophan levels in serum, urine and feces samples of age-matched monkeys with or without pre-diabetes and diabetes (n=26/group). (E-F) Correlation analysis between fecal tryptamine level and fasting blood glucose (FBG) and HbA1c in monkeys with or without pre-diabetes and diabetes (n=26/group). (G) Tryptamine production in batch culture experiments using feces from monkeys age-matched monkeys with or without pre-diabetes and diabetes (n=26/group). (H-I) Tryptamine and tryptamine/tryptophan ratio in fecal samples of human with or without type 2 diabetes (n=25 healthy controls, n=15 diabetic patients). (J) Spearman correlation of fecal tryptamine/tryptophan ratio with blood glucose level in human with or without type 2 diabetes (n=25 healthy controls, n=15 diabetic patients). Data were presented as mean ± S.D. *P*-values were determined by ordinary one-way ANOVA, Student’s t-test and spearman’s rank correlation. See also Figure.S1.

We then performed a targeted metabolomics analysis to quantify host and gut-microbial metabolites of tryptophan in serum, urine and feces of diabetic monkeys. Notably, tryptamine, a microbial metabolite transformed from tryptophan by microbial tryptophan decarboxylase in gut, is found significantly increased in feces of diabetic monkeys (Figure.1C). By contrast, tryptophan (Trp, the precursor of tryptamine) level is not found different in feces, suggesting increased tryptamine may be attributed to the increased catabolism of tryptophan by microbial tryptophan decarboxylase but not tryptophan level in diabetes (Figure.1D). Correlation analysis between fecal tryptamine level and T2D-related blood characteristics in monkeys revealed tryptamine is positively correlated with fasting blood glucose (FBG) and Hb1Ac in diabetic monkeys (Figure.1E-F). Moreover, we cultured fecal bacterial suspension from fecal samples of each monkey supplemented with tryptophan under anaerobic conditions. A higher concentration of tryptamine was found in the culture medium from the diabetic group, indicating diabetes-associated microbiota has a higher catalytic ability to transform tryptophan into tryptamine (Figure.1G).

To further validate the clinical relevance of tryptamine in T2D, we determined fecal tryptamine level in healthy controls and T2D subjects recruited from a published study (Zheng et al., 2021). In line with our findings in spontaneous diabetic monkeys, both fecal tryptamine and its ratio with tryptophan significantly increased in diabetic patients compared with healthy volunteers (Figure.1H-I). Moreover, correlation analysis showed fecal tryptamine is positively correlated with FBG in T2D patients (Figure.1J). Together, these results showed tryptamine level is increased in both diabetic monkeys and T2D patients, and positively correlated with blood glucose indexes, revealing the potential detrimental role of tryptamine in the development of T2D.

### Dietary fibers intervention suppresses tryptamine level in subjects with type 2 diabetes

We have previously reported a dietary fiber intervention improves glucose intolerance in T2D subjects (Zhao et al., 2018). We therefore determined tryptophan and tryptamine level in fecal samples of T2D subjects before and after treatment of dietary fiber, in order to study whether dietary fiber intervention can suppress tryptamine level and correlation between tryptamine and glucose tolerance in this study. Interestingly, fecal tryptamine level and tryptamine/tryptophan ratio were found significantly suppressed in dietary fiber-treated subjects (W group) but not in control group subjects (U group) after 12 weeks treatment (Figure.2A-B). Tryptamine level was also found negatively correlated with improvement of glucose tolerance indexes including HbA1c, OGTT and homeostatic model assessment for insulin resistance (HOMA-IR) indexes in W group but not in U group (Figure.2C-E and Supplement Figur.S2A-C). These findings validated tryptamine level is positively correlated with glucose tolerance, suggesting tryptamine is a potential pathogenic factor of T2D through manipulation of glucose tolerance.

**Figure.2.**
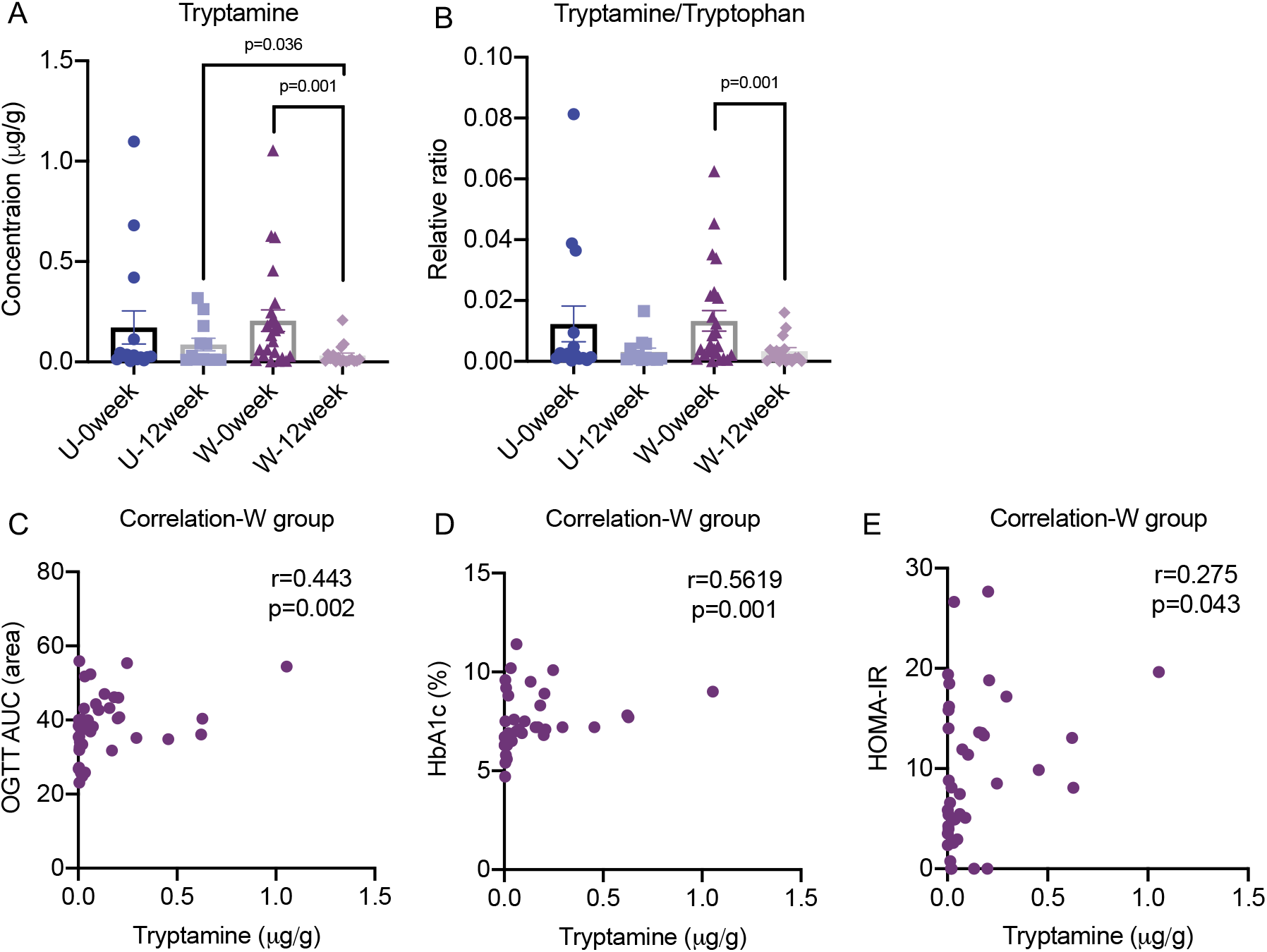
Dietary fibers intervention suppresses tryptamine level in T2D subjects. (A-B) Tryptamine and tryptamine/tryptophan ratio in fecal samples of control group (U group; n=16) and high-fiber group (W group; n=27) of T2D subjects (Day 0 and Day 84). (C-E) Correlation analysis between fecal tryptamine level and OGTT, HbA1c and HOMA-IR indexes in T2D subjects treated with high-fiber (W group; n=27). *P*-values were determined by one-way ANOVA, paired t-test and spearman’s rank correlation. See also Figure.S2

### Tryptamine impairs glucose tolerance and insulin sensitivity

Since we have showed that tryptamine is positively associated with glucose tolerance in diabetic monkeys and T2D subjects, we wondered whether tryptamine impairs glucose tolerance. Based on the tryptamine concentration we determined in fecal samples of T2D subjects, we studied tryptamine action at a dosage of 10mg/kg on glucose tolerance in normal monkeys using intravenous glucose tolerance test (IVGTT). Notably, tryptamine-treated monkeys had higher glucose and insulin level (Figure.3A-B), suggesting tryptamine treatment significantly induce glucose intolerance. Following this finding, we examined tryptamine-treated mice by oral glucose tolerance test (OGTT) experiments to further validate whether tryptamine impairs glucose tolerance. As expected, OGTT showed the tryptamine-treated mice had higher blood glucose levels starting at a dosage of 2 mg/kg, suggesting tryptamine impairs glucose tolerance in normal mice (Figure.3C). We also found a significant elevation in serum insulin level of tryptamine-treated mice compared with control mice (Figure.3D). In line with the effect of acute treatment of tryptamine on glucose tolerance, long-term treatment of tryptamine also impairs glucose tolerance in HFD-fed mice (Supplement Figure. S3A). These results suggest tryptamine impairs glucose tolerance and stimulates insulin secretion, thereafter tryptamine may inhibit insulin sensitivity to impair the glucose tolerance.

**Figure.3.**
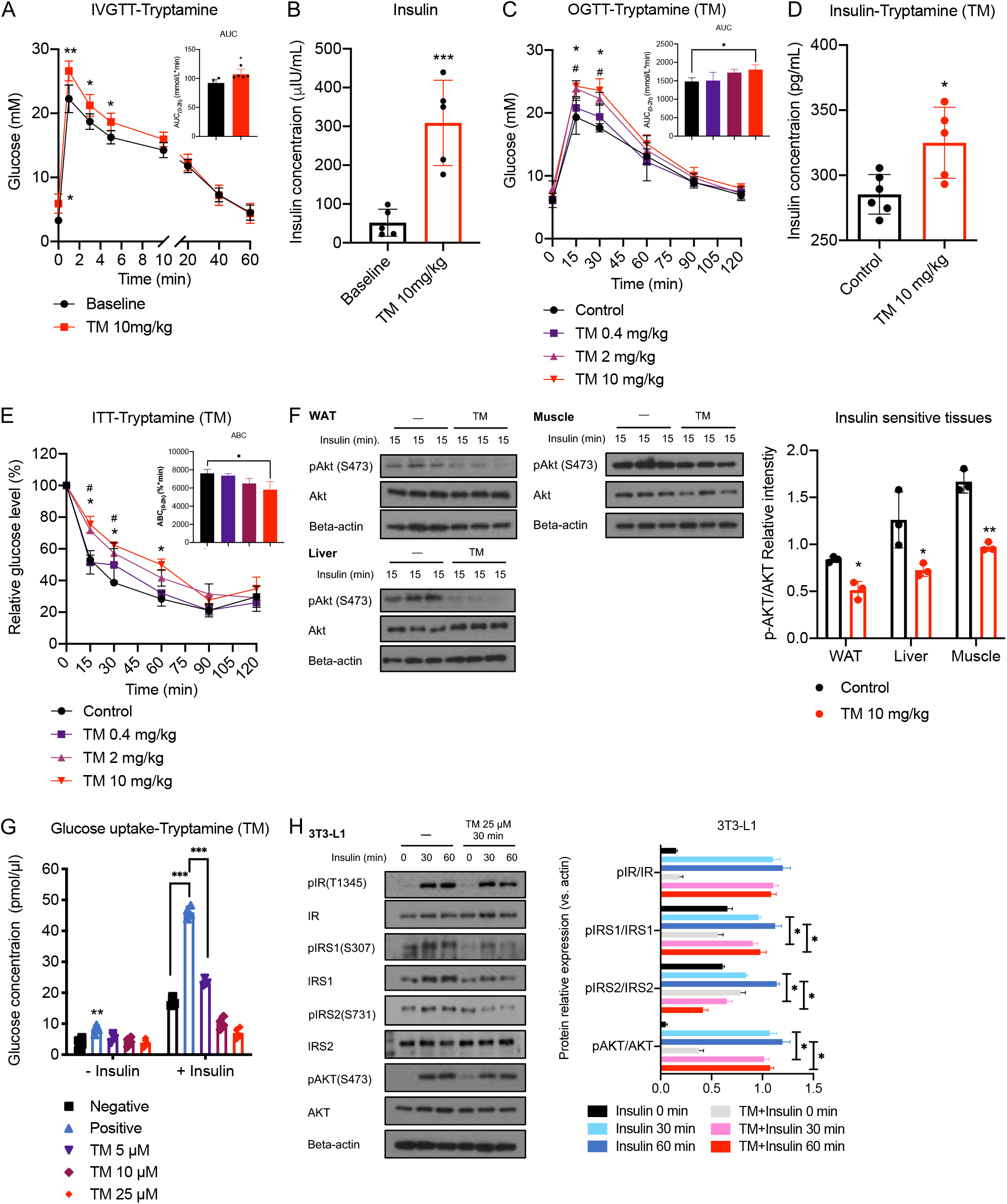
Tryptamine impairs glucose tolerance and insulin sensitivity in monkeys and mice. (A) Intravenous glucose tolerance test (IVGTT) in monkeys after treatment of tryptamine (10mg/kg) or control (1% CMC-Na in water) (n=6/group). (B) Insulin level in serum in monkeys after treatment of tryptamine (10mg/kg) or control (1% CMC-Na in water) (n=6/group). (C/E) Oral glucose tolerance test (OGTT) and insulin tolerance test (ITT) in normal mice after treatment of tryptamine as indicated dosages (0.4mg/kg, 2mg/kg and 10mg/kg) or control (1% DMSO in saline) (n=6/group). * comparisons between control group and tryptamine group (10 mg/kg). # comparisons between control group and tryptamine treatment (2 mg/kg). (D) Insulin level in serum in mice after treatment of tryptamine (10mg/kg) or control (1% DMSO in saline) (n=6/group). (F) Western blot (and semi-quantification) of tryptamine treatment (10mg/kg) on Akt activation stimulated by insulin (1U/kg) in white adipose tissue (WAT) lysates, liver lysates and skeletal muscle lysates from mice. (n=3/group) (G) Effect of the tryptamine (5μM, 10μM and 25μM) on glucose uptake stimulated by insulin (10nM) in 3T3-L1 cells (n=3/group). (H) Western blot (and semi-quantification) of tryptamine treatment (25μM) on insulin signaling stimulated by insulin (10nM) in 3T3-L1 cells (n=3/group). Data were presented as mean ± S.D. *, p < 0.05, **, p < 0.01 and ***, p < 0.001. *P* values were determined by ordinary one-way ANOVA or Student’s t-test. See also Figure. S3.

We then conducted the insulin tolerance test (ITT) to determine whether tryptamine affects insulin sensitivity in mice. ITT experiment showed tryptamine-treated mice had higher glucose levels at a dosage of 2mg/kg (Figure.3E). Following that, we examined the tryptamine effects on AKT phosphorylation in mice. In line with ITT result, tryptamine treatment significantly downregulated AKT phosphorylation in insulin-stimulated insulin-sensitive tissues including white adipose, liver and skeletal muscle, suggesting acute tryptamine treatment impairs insulin sensitivity *in vivo* (Figure.3F). Besides *in vivo* studies, we also determined the tryptamine effects on glucose uptake and key components of insulin signaling in 3T3-L1 adipocytes. Tryptamine treatment inhibited basal glucose uptake in a dose-dependent manner (Figure.3G) and insulin signaling in a time-dependent manner in 3T3-L1 cells (Figure.3H), indicating tryptamine impairs glucose uptake and insulin sensitivity *in vitro.*

Tryptamine can be rapidly metabolized by the host monoamine oxidase into indole-3-acetic acid (IAA) within 30 min after entering the circulation from the GI tract (Modoux et al., 2021). To investigate whether gut microbiota-derived tryptamine can enter the insulin-sensitive tissues, we gave mice tryptamine via oral gavage to mimic the increased fecal tryptamine level in diabetic monkeys and T2D subjects. After oral administration, tryptamine and IAA levels were significantly increased in serum and insulin-sensitive tissues including WAT, liver and muscle within 15 min (Supplement Figure.3B-C), indicating tryptamine can rapidly enter insulin-sensitive tissues and the inhibitory effects of tryptamine on insulin sensitivity is possibly mediated by tryptamine or its metabolite IAA. We then studied whether the precursor (Trp) or metabolite (IAA) of tryptamine can affect glucose tolerance or insulin sensitivity. Trp has been shown to suppress glucose level and preserves insulin secretion *in vivo* (Goodarzi et al., 2021; Inubushi et al., 2012), and no significant difference in blood glucose level between the control group and IAA-treated group was found in the OGTT experiment (Supplement Figure. S3D). Moreover, Trp and IAA treatment did not affect the AKT phosphorylation in insulin-stimulated 3T3-L1 cells (Supplement Figure.S3E), suggesting tryptamine but not its precursor or metabolite can impair glucose tolerance and insulin sensitivity. Together, these *in vivo* and *in vitro* data showed tryptamine impairs glucose tolerance and insulin sensitivity.

### Tryptamine weakens insulin signaling via TAAR1-ERK signaling axis

To investigate the underlying mechanism of tryptamine effects on insulin sensitivity, we used phospho-proteomics approach to determine the systemic changes in phosphorylation sites of proteins with and without tryptamine treatment in WAT of normal mice. Through phospho-proteomics analysis and KEGG mapping, we found insulin signaling-related proteins namely hormone-sensitive lipase (HSL), mitogen-activated protein kinase MAPK 1/3 (ERK), sorbin and SH3 domain containing 1 (SH3D5) are upregulated by tryptamine treatment (Supplement Table S3-S4). Among these proteins, MAPK/ERK has been implicated in the development of insulin resistance in T2D and its phosphorylation levels have been found to increase in adipose tissue of animal models of T2D and T2D patients (Ozaki et al., 2016). Therefore, we speculated gut bacteria-derived tryptamine may activate ERK pathway to suppress insulin sensitivity in T2D.

We firstly validated the ERK phosphorylation levels in tryptamine-treated mice and found p-ERK1/2 is upregulated in insulin-sensitive tissues (Figure.4A). Next, we used ERK inhibitors (U0126 and PD98059) to determine whether the suppressing action of tryptamine on glucose tolerance and insulin sensitivity is dependent on MAPK/ERK pathway. Treatment of ERK inhibitors significantly improved glucose tolerance and insulin sensitivity in mice treated with tryptamine (Figure.4B-C and Supplement Figure. S4A-B) in OGTT and ITT study. In contrast, ERK inhibitors did not affect glucose tolerance or insulin sensitivity in control mice.

**Figure.4.**
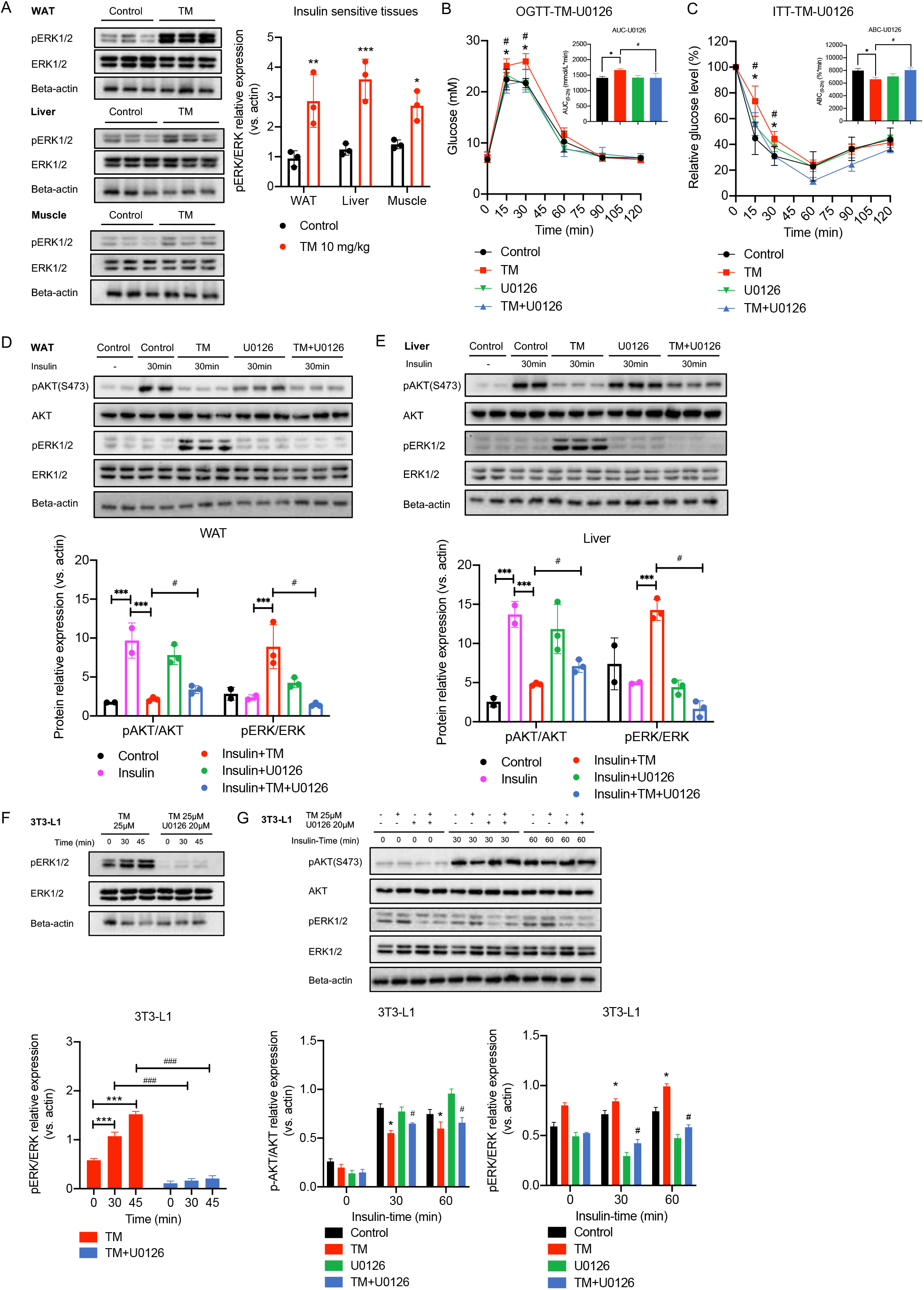
Tryptamine suppresses insulin signaling via ERK activation. (A) Western blot (and semi-quantification) of tryptamine treatment (10mg/kg) on ERK activation in white adipose tissue (WAT) lysates, liver lysates and skeletal muscle lysates from mice (n=3/group). (B-C) Oral glucose tolerance test (OGTT) and insulin tolerance test (ITT) in mice after treatment of tryptamine (10mg/kg), ERK inhibitor U0126 (20mg/kg) or control (1% DMSO in saline) (n=6/group). * comparisons between control group and tryptamine group (10 mg/kg). # comparisons between tryptamine group and tryptamine + ERK inhibitor (U0126) group. (D-E) Western blot (and semi-quantification) of tryptamine (10mg/kg) and ERK inhibitor U0126 treatment (20mg/kg) on ERK activation and Akt activation stimulated by insulin (1U/kg) in white adipose tissue (WAT) lysates and liver lysates from mice (n=2-3/group). (F) Western blot (and semi-quantification) of tryptamine treatment (25μM) and ERK inhibitor U0126 on ERK activation in 3T3-L1 cells (n=3/group). (G) Western blot (and semi-quantification) of tryptamine (25μM) and ERK inhibitor U0126 treatment (20μM) on ERK activation and Akt activation stimulated by insulin (10nM) in 3T3-L1 cells (n=3/group). Data were presented as mean ± S.D. *, # p < 0.05, **, ## p < 0.01 and ***, ### p < 0.001. *P* values were determined by ordinary one-way ANOVA or Student’s t-test. See also Figure.S4.

Consistently, ERK inhibitors abolished the inhibitory effects of tryptamine on the insulin-stimulated AKT phosphorylation in insulin-sensitive tissues and significantly downregulated tryptamine-induced ERK phosphorylation (Figure.4D-E). In line with *in vivo* observation, we showed that tryptamine treatment significantly upregulated ERK phosphorylation in a time-dependent manner in 3T3-L1 cells, which was blocked by pretreatment of ERK inhibitors (Figure.4F and Supplement Figure.S4C). ERK inhibitors significantly suppressed the inhibitory effects of tryptamine on AKT phosphorylation in insulin-stimulated 3T3-L1 cells, suggesting tryptamine impairs insulin sensitivity via activating MAPK/ERK pathway (Figure.4G and Supplement Figure.S4D).

Tryptamine can bind to GPCR receptor TAAR1, which exhibits stimulatory effects for the downstream MAPK/ERK pathway (Zucchi et al., 2006). We speculated gut bacteria-derived tryptamine may activate TAAR1/MAPK/ERK pathway to inhibit insulin sensitivity in T2D so we investigated whether TAAR1 antagonism can protect against tryptamine-induced impairment of glucose tolerance and insulin sensitivity via the MAPK/ERK pathway. In the OGTT and ITT study, treatment with EPPTB, a specific TAAR1 antagonist, ameliorated the effect of tryptamine on glucose tolerance and insulin sensitivity (Figure.5A-B). EPPTB treatment also significantly downregulated tryptamine-induced ERK phosphorylation and the inhibitory effects of tryptamine on insulin-stimulated AKT phosphorylation were abolished by EPPTB (Figure.5C-D). By contrast, EPPTB did not affect glucose tolerance or insulin sensitivity in control mice. Moreover, we examined the inhibitory effects of EPPTB on the cAMP level and phosphorylation level of ERK in tryptamine-treated 3T3-L1 cells. In line with *in vivo* results, EPPTB treatment significantly downregulated the increased cAMP level and ERK phosphorylation induced by tryptamine and antagonized the inhibitory effects of tryptamine on AKT phosphorylation in 3T3-L1 cells (Figure.5E-G). Several receptors of tryptamine including TAAR1, AhR and 5-HTR4 have been identified before (Cheng et al., 2015). In the present study, we found tryptamine activation on ERK can be suppressed by TAAR1 antagonist EPPTB but not AhR antagonist or 5HT antagonists in 3T3-L1 cells, suggesting TAAR1 plays an important role in mediating the tryptamine action on ERK signaling and insulin signaling (Supplement Figure. S5A-C). In summary, we showed tryptamine weakens insulin signaling via TAAR1-ERK signaling axis, results in impaired glucose tolerance and insulin sensitivity.

**Figure.5.**
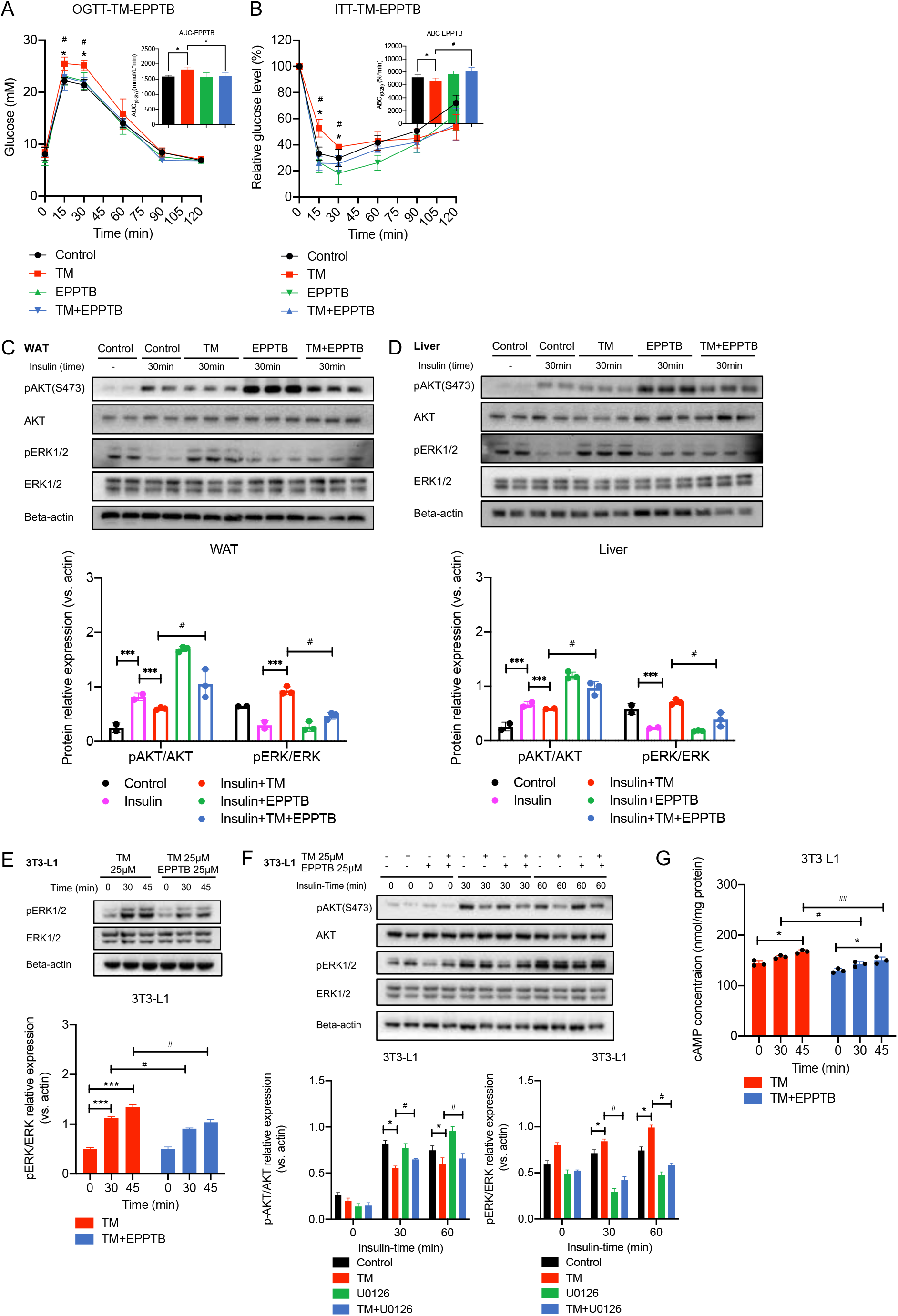
Tryptamine impairs insulin signaling via TAAR1-ERK signaling axis. (A-B) Oral glucose tolerance test (OGTT) and insulin tolerance test (ITT) in mice after treatment of tryptamine (10mg/kg), TAAR1 antagonist EPPTB (10mg/kg) or control (1% DMSO in saline) (n=6/group). * comparisons between control group and tryptamine group (10 mg/kg). # comparisons between tryptamine group and tryptamine + TAAR1 antagonist (EPPTB) group. (C-D) Western blot (and semi-quantification) of tryptamine (10mg/kg) and TAAR1 antagonist EPPTB (10mg/kg) treatment on ERK activation and Akt activation simulated by insulin (1U/kg) in white adipose tissue (WAT) lysates and liver lysates from mice (n=2-3/group). (E) Western blot (and semi-quantification) of tryptamine (25μM) and TAAR1 antagonist EPPTB treatment (20μM) on ERK activation in 3T3-L1 cells (n=3/group). (F) Western blot (and semi-quantification) of tryptamine (25μM) and TAAR1 antagonist EPPTB treatment (20μM) on ERK activation and Akt activation simulated by insulin (10nM) in 3T3-L1 cells (n=3/group). (G) cAMP measurement of tryptamine (25μM) and TAAR1 antagonist EPPTB (20μM) treatment in 3T3-L1 cells (n=3/group). Data was presented as mean ± S.D. *, # p < 0.05, **,## p < 0.01 and ***, ### p < 0.001. *P* values were determined by ordinary one-way ANOVA or Student’s t-test. See also Figure.S5.

### *In vivo* tryptamine production by type 2 diabetes-associated tryptamine-producing bacteria impairs insulin sensitivity

To understand the role of tryptamine-producing bacteria in the development of T2D, we identified tryptamine-producing bacteria in our T2D dataset (GUT2D) (Zhao et al., 2018) based on two reference tryptophan decarboxylase (TDC) sequences (Williams et al., 2014). Notably, a total of 5 bacterial co-abundance groups (CAGs) namely *Ruminococcus gnavus* CAG0075, *Klebsiella* CAG0098, *Alistipes putredinis* CAG0116, *Rikenellaceae bacterium* CAG0092 and *Bacteroides caccae* CAG0012 were found with a prevalence above 20% in gut metagenome in GUT2D. Among them, *Klebsiella* CAG0098 was a non-responder while the other four CAGs were negative responders of dietary intervention (Supplement Table. S5). Since we have shown tryptamine is suppressed by dietary intervention in this study, we performed correlation analysis between tryptamine and these gut bacteria to identify T2D-associated tryptamine producers. Interestingly, correlation analysis showed only *R. gnavus* CAG0075 is positively correlated with tryptamine and tryptamine/tryptophan ratio among 5 CAGs (Supplement Table.S6). Considering TDC sequence belonged to *R. gnavus* CAG0075 had 100% identity compared with the reference sequence from *R. gnavus* (ATCC 29149) and the average nucleotide identity between *R. gnavus* CAG0075 and *R. gnavus* (ATCC 29149) was 99.1% (Supplement Table. S5), we demonstrated *R. gnavus* (ATCC 29149) is a T2D-associated tryptamine-producing bacteria strain.

We then performed monoassociation study in pseudo germ-free mice to investigate *R. gnavus* (ATCC 29149) action on glucose tolerance and insulin signaling. TAAR1 antagonist EPPTB was used to block tryptamine action on insulin signaling. Notably, fecal tryptamine level in pseudo germ-free mice was found significantly reduced after receiving an antibiotics cocktail (Figure.6A), and colonization with *R. gnavus* significantly upregulated fecal tryptamine level (Figure. 6D and Supplement Figure. S6A). In line with previous findings, impaired OGTT, ITT and AKT phosphorylation indexes were observed in mice colonized with *R. gnavus* and partially abolished by EPPTB treatment (Figure.6B-E). These results demonstrated monoassociation of T2D-associated tryptamine producers (eg. *R. gnavus*) impaired insulin sensitivity, contributing to the development of insulin resistance in T2D.

**Figure.6.**
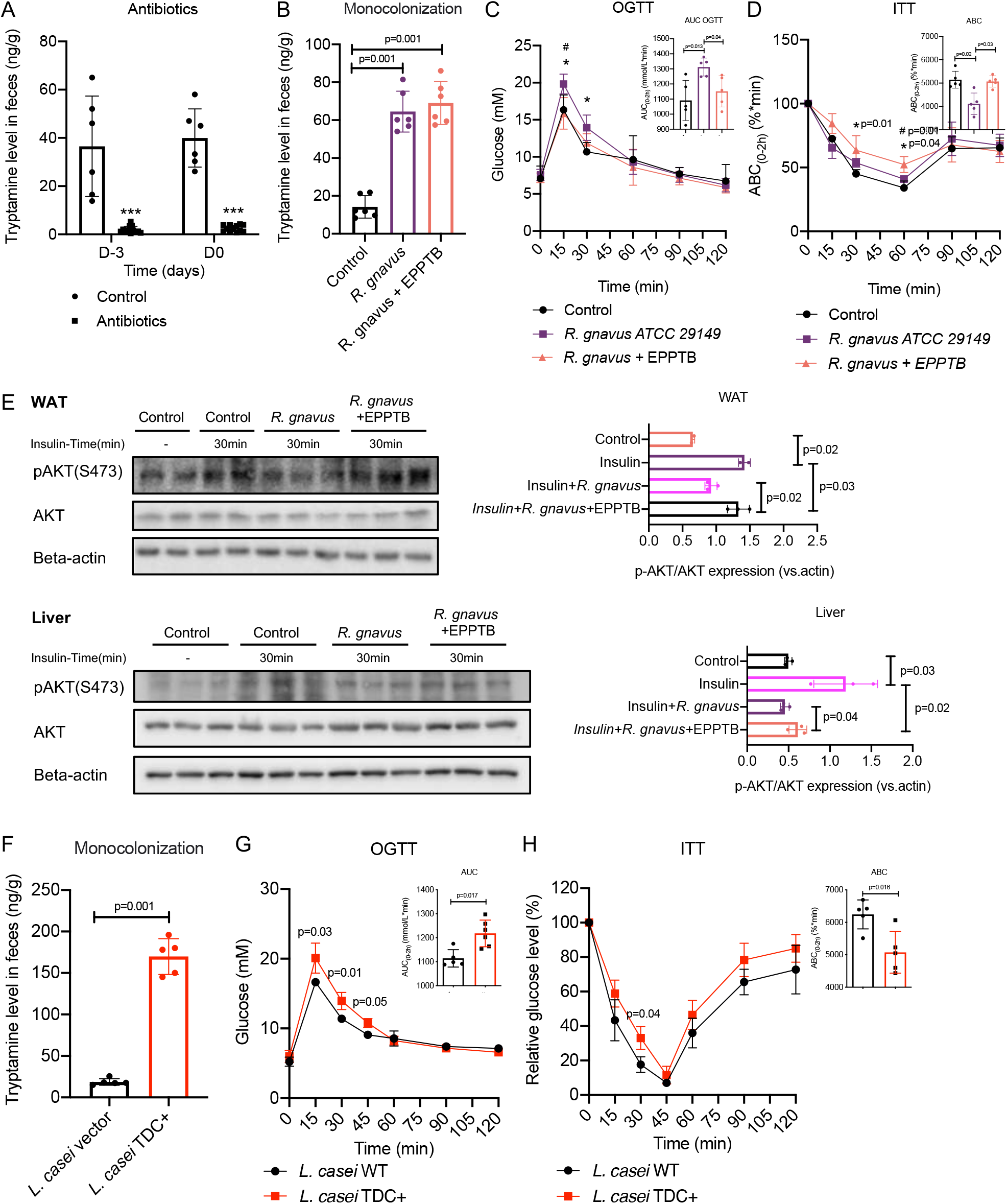
*In vivo* tryptamine production by T2D-associated tryptamine producers and engineered gut microbe impairs insulin signaling. (A) Fecal tryptamine level in mice from control group (n=6) and antibiotics mixture treatment (ABX) group (n=12). (B-D) Fecal tryptamine level, oral glucose tolerance test (OGTT) and insulin tolerance test (ITT) indexes in antibiotics mixture (ABX)-treated mice following colonization of either *R. gnavus* ATCC 29149 and i.p. administration of TAAR1 antagonist EPPTB (10mg/kg) (n=6 per group). * comparisons between control group and *R. gnavus* group. # comparisons between *R. gnavus* group and *R. gnavus* group + TAAR1 antagonist (EPPTB) group. (E) Western blot (and semi-quantification) of *R. gnavus* ATCC 29149 and EPPTB administration treatment on Akt activation simulated by insulin (1U/kg) in white adipose tissue (WAT) lysates from pseudo germ-free mice (n=2-3 per group). (F-H) Fecal tryptamine level, oral glucose tolerance test (OGTT) and insulin tolerance test (ITT) indexes in antibiotics mixture (ABX)-treated mice following colonization of either engineered *L. casei* TDC+ or *L. casei* with empty vector (n=6 per group). (I) Western blot (and semi-quantification) of colonization of either engineered *L. casei* TDC+ or *L. casei* with empty vector on Akt activation simulated by insulin (1U/kg) in white adipose tissue (WAT) lysates from pseudo germ-free mice (n=2-3 per group). *, # p < 0.05, **,## p < 0.01 and ***, ### p < 0.001. *P* values were determined by ordinary one-way ANOVA or Student’s t-test. See also Figure.S6.

To testify whether *in vivo* tryptamine production by gut bacterial TDC impair glucose tolerance and insulin sensitivity, we integrated the plasmid expressing TDC from *R*. *gnavus* (ATCC 29149) into a wild type gut microbe *Lactobacillus casei.* We confirmed that tryptamine level was significantly increased in culture MRS medium (supplemented with 0.25% Trp) from engineered *L. casei* TDC^*+*^ strain compared to the *L. casei* with empty vector insertion and medium control after overnight inoculation (Supplement Figure.S6B). Next, to investigate whether the engineered *L. casei* TDC^*+*^ strain enhance fecal tryptamine level, we colonized the pseudo germ-free mice with *L. casei* TDC^*+*^ or blank vector *L. casei.* Notably, pseudo germ-free mice colonized with *L. casei* TDC^*+*^ exhibited increased levels of fecal tryptamine in feces compared with blank vector *L. casei,* confirming that *L. casei* TDC^*+*^ produced tryptamine *in vivo* (Figure.6F). Accompanied with the increased fecal tryptamine level, higher OGTT, ITT indexes and impaired AKT phosphorylation were found increased in pseudo germ-free mice colonized with *L. casei* TDC^*+*^ compared with blank vector *L. casei* (Figure.6G-I), indicating *in vivo* gut bacteria TDC enhanced fecal tryptamine level and impaired insulin sensitivity, thus contributing to the insulin resistance in T2D patients.

## Discussion

In the present study, we provided novel mechanistic insights into the contribution of gut microbiota and its metabolite tryptamine to the development of T2D. We showed that tryptamine impaired glucose tolerance and insulin sensitivity. Tryptamine production was increased in monkeys and human with T2D and it was positively correlated with key glucose homeostasis parameters including FBG and Hb1Ac. We also showed tryptamine downregulation is positively correlated with improvement of glucose tolerance in a dietary fiber intervention study. Mechanistically, tryptamine inhibits insulin sensitivity via the TAAR1-ERK signaling axis. The clinical relevance of T2D-associated tryptamine producers in host glucose homeostasis is further validated by monoassociation of *R. gnavus* and engineered *L. casei* to produce tryptamine in pseudo GF mice. In summary, this study showed gut microbiota-derived tryptamine is positively associated with the development of T2D and contributes to the insulin resistance by impairing insulin signaling via TAAR1-ERK signaling axis. Tryptamine is a newly characterized pathogenic factor for T2D and may serve as a therapeutic target for the treatment of T2D.

Despite the fact that gut microbiome composition and structure vary considerably across geographic locations, races and ethnicities, the differences in the microbial metabolites profiles are much smaller than that observed in taxonomy (Visconti et al., 2019), therefore investigation of gut microbial metabolites can help reveal the role of gut microbiota in the pathogenesis of T2D. In this study, normal and diabetic crab-eating macaques were chosen to study the changes of gut microbial metabolites altogether with healthy controls and T2D subjects. The diabetic crab-eating macaques were used in the metabolomics study for several reasons. First, they spontaneously develop similar metabolic phenotypes of T2D patients including hyperglycemia and insulin resistance. Second, they share a similar structure and composition of gut microbiota as human compared with rodent animals, and they are less affected by confounding factors that can affect the gut microbiome in human gut microbiome study such as age, anti-diabetic medication and dietary pattern (Chen et al., 2018; Nagpal et al., 2018). Third, the fecal metabolome in diabetic crab-eating macaques has not been studied before, thereafter our investigation can provide new understanding towards the metabolic profiles of pre-clinical T2D models. In addition to tryptamine elevation, a series of metabolites derived from bile acid metabolism, amino acids metabolism and purine metabolism are also found significantly changed in fecal metabolome of spontaneous diabetic monkeys (Supplement Table.S1). Collectively, this model is applicable for studying the changes of gut-microbial metabolites in T2D.

Tryptamine as a microbial metabolite, has shown multiple biological activities including beneficial and detrimental effects to host. Recent studies reported tryptamine is increased in patients with diarrhea-dominant irritable bowel syndrome (Mars et al., 2020) and accelerates gastrointestinal transit by activating 5-HT4 receptor (Bhattarai et al., 2018). Gut microbiota-derived tryptamine improves intestinal barrier integrity and barrier functions via activating aryl hydrocarbon receptor (AhR) (Scott et al., 2020) and stimulates mucus release from goblet cells via 5-HTR4 agonism (Bhattarai et al., 2020). Tryptamine possess pharmacological effects on attenuating experimental multiple sclerosis in mice via activating AhR (Dopkins et al., 2021). Similarly, tryptamine attenuated pro-inflammatory responses in murine macrophages and hepatocytes in a AhR dependent mechanism (Krishnan et al., 2018). Our study revealed tryptamine contributes to the development of T2D by impairing insulins sensitivity. These research findings suggest gut microbiota-derived tryptamine is considerably involved in the host health via host-microbiota interactions and may also play pivotal roles in other diseases.

After showing tryptamine impairs insulins sensitivity, we revealed tryptamine induces a significant activation of ERK signaling pathway in insulin sensitive tissues. ERK activity is elevated in WAT of humans and rodents in diabetic conditions (Bashan et al., 2007; Carlson et al., 2003). Activation of the MEK1-ERK pathway significantly suppressed the protein expression of IR, IRS-1 and IRS-2 as well as the insulin-stimulated phosphorylation of IR, IRS-1 and IRS-2, and the downstream PI3K-AKT signaling pathway (Fujishiro et al., 2003). Moreover, specific inhibition of the ERK pathway using chemical inhibitors is effective to improve insulin resistance in *db/db* and HFD-fed mice (Ozaki et al., 2016), representing a potential therapeutic strategy to combat tryptamine-induced insulin resistance.

We further demonstrated tryptamine action on ERK activation and inhibition of insulin signaling pathway can be abolished by TAAR1 antagonist. Tryptamine is a ligand of several receptors including TAAR1 (Miller, 2011), AhR (Vikström Bergander et al., 2012) and serotonin receptors (Yasi et al., 2019). Although AhR and serotonin receptors are extensively studied for their important role in metabolic regulation (Kim et al., 2013; Oh et al., 2016; Wang et al., 2011), we showed tryptamine activation on ERK can be suppressed by TAAR1 antagonist but not AhR antagonist or 5HT antagonists. Further studies are required to uncover whether these receptors are also participated in the tryptamine action on insulin sensitivity through other signaling mechanisms.

TAAR1 is found expressed in both central nervous system and gut (Sotnikova et al., 2009). TAAR1 activator has recently been proposed as a potential target for improving glycemic control and TAAR1/Gas-mediated signaling pathways also promote insulin secretion (Michael et al., 2019). In contrast, our study showed long-term exposure to TAAR1 activator tryptamine impairs insulin sensitivity, thereafter the pharmacological use of TAAR1 modulator on glucose control requires carefully studied. TAAR1 can also be activated with aromatic amino acid-derived biogenic amines from gut microbes such as tyramine and beta-phenethylamine (Zucchi et al., 2006). Considering tryptamine as microbial TAAR1 ligands impairs insulin sensitivity, further studies with tyramine and beta-phenethylamine are needed be addressed to gain a more comprehensive understanding of the action of amino acid-derived biogenic amines on T2D. Interestingly, a significant elevation of fecal beta-phenethylamine but not tyramine was found increased in diabetic monkeys and T2D patients (Supplement Figure. S1F-I), therefore beta-phenethylamine may also be involved in the development of insulin resistance and pathogenies of T2D, which deserves further investigations.

Besides of the mechanistic insights into the contribution of gut microbiota-derived tryptamine to the development of T2D, we are the first to report T2D-associated tryptamine producers (*R. gnavus*) impairs insulin sensitivity, thus contributing to the development of insulin resistance. Tryptamine-producing bacteria including *Ruminococcus gnavus* and *Clostridium sporogenes* have been identified and a variety of gut bacteria including *Lachnospiraceae bacterium* 2_1_58FAA, *Blautia hansenii, Clostridium nexile, Desulfitobacterium dehalogenans*, and *Clostridium asparagiforme* were also found highly similar to TDC gene of *R. gnavus* (RUMGNA_01526) by a previous study (Williams et al., 2014). Considering literature has provided several bacteria species can potentially produce tryptamine, we also characterized tryptamine-producing bacteria in human gut metagenome based on TDC sequences using a computational approach. A total of 37 gut bacteria species that potentially capable of producing tryptamine were identified (Supplement Table.S7) and some bacteria species were also found significantly increased in monkey and human with T2D, supporting the increased tryptamine production is attributed to the increased abundances of those T2D-associated gut bacteria (Supplement Figure.S7). Due to the lack of commercially available bacteria strains, the experimental validation of these potential tryptamine-producing bacteria and their effects on glucose tolerance will be studied in future.

Collectively, our data suggest gut microbiota-derived tryptamine impairs insulin sensitivity via a TAAR1-ERK signaling axis, thereby contributing to the development of T2D. Our findings not only provide new insights into the role of gut microbiota in the pathogenesis of T2D, but also construct fundamental knowledge for developing gut microbiota-based therapeutic targets for management of T2D.

## Acknowledgments

This work was kindly funded by the National Natural Science Foundation of China (81973538); Shenzhen Science and Technology Innovation Committee (JCYJ20190808164201654); Medical Science and Technology Research Foundation of Guangdong Province (A2020272) and Key-Area Research and Development Program of Guangdong Province (2020B1111110003). We are thankful to all patients and healthy volunteers who donated specimens for this study.

## Author contributions

W.J., LP.Z. and ZX.B. designed the experiments and revised the manuscript in this study. LX.Z. and HT.X. wrote the manuscript with input from co-authors. LX.Z., CY. L., MX. G. and ZW.N. conduced metabolomics study in monkeys and human. LX.Z., HT.X. and CY.L. performed mice experiments. GJ.W, YS. D. and C.Y. performed metagenomics and computational analysis. LX.Z., CH.H., YJ.Z. and M.Z. conduced the cell and bacteria experiments. EL. Z., L.Z., HLX. W., YY.L., CH.Z. and XD.F. contributed to the study design, technical support and data analysis support towards this work. W.J, CH.Z. and LP.Z. provided clinical recourses and contributed to the study design.

## Declaration of interests

The authors have claimed no financial interests to declare.

## STAR Methods

### Contact for reagent and resource sharing

Further information and requests for resources and reagents should be directed to and will be fulfilled by the Lead Contact Zhao-xiang Bian.

### Experimental models and subject details

#### Monkey study

The first crab-eating macaques (*Macaca fascicularis*) metabolomics study is performed with ethics approval (No. YMB1704) by Yunnan Yinmore Biotechnology company (Kunming, China). The diagnosis of diabetes for monkeys were based on publish criteria (Auerbach et al., 2006; Harwood Jr et al., 2012; Vaughan and Mattison, 2016). Specifically, age-matched (10-23 years old) monkeys were categorized as normal (Define FBG <75mg/dL and Hb1Ac <3.5%), pre-diabetes (FBG 80-130 mg/dL and Hb1Ac 4.0%-6.0%) and diabetes (FBG >130 mg/dL and Hb1Ac > 6.0 %) (n=26 for each group). Biological samples including serum, urine and feces were collected and stored at −80°C until analysis. The second crab-eating macaques (n=5) intervention study is performed with ethics approval (No. HZ2021047) by Huazhen Biosciences company (Guangzhou, China). Biochemical parameters including OGTT and serum insulin level were measured before and after tryptamine treatment. No crab-eating macaques were treated with anti-diabetic medication during these studies.

#### Mouse study

All mouse studies were approved by the Committee on the Use of Human & Animal Subjects in Teaching & Research at Hong Kong Baptist University (Hong Kong SAR, China) and performed following the Animals (Control of Experiments) Ordinance of Department of Health, Hong Kong SAR, China and reported following the ARRIVE guidelines (du Sert et al., 2020). Male C57BL/6J mice aged 6-8 weeks and weighed 20-25g were purchased from Laboratory Animal Services Centre, The Chinese University of Hong Kong (Hong Kong SAR, China). The mice were housed with a 12 h light/dark cycle at a controlled temperature of around 25°C with free access to food and water.

Pseudo germ-free mice were generated using antibiotics cocktails containing 50mg/kg vancomycin, 100mg/kg neomycin, 100mg/kg metronidazole, 100mg/kg ampicillin, 50mg/kg streptomycin via oral gavage for 9 days (one time per day) as previously reported (Zhao et al., 2019). Fecal samples were collected as required for the tryptamine measurement.

#### Human study

The first cohort (Zheng et al., 2021) of healthy controls and T2D subjects was approved by the Research Ethics Committee of Shanghai Jiao Tong University Affiliated Sixth People’s Hospital. Written informed consent was signed and obtained from all participants. Subjects with fasting blood glucose < 6.1 mmol/L were classified as healthy controls (HC), whereas those with fasting blood glucose > 7.0 mmol/L or OGTT (2h) > 11.1 mmol/L were classified as T2D.

The second cohort GUT2D study (Zhao et al., 2018) was approved by the Ethics Committee at School of Life Sciences and Biotechnology, Shanghai Jiao Tong University (ID: 2014-016). Written informed consent was obtained from all participants. The trial was registered in the Chinese Clinical Trial Registry (ChiCTR-TRC-14004959). Participants were received either acarbose plus usual diet (control; U group) or acarbose plus the WTP diet (intervention; W group) for 84 days.

#### Cell study

3T3-L1 adipocytes (ATCC CL-173) were cultured and maintained in DMEM with 10% (v/v) FBS. For the glucose uptake assay, 3T3-L1 cells were serum- and glucose-starved for 3 hours and then incubated with glucose, FBS, insulin and tryptamine for 30 min as indicated. To assess the effect of tryptamine on insulin signaling, 3T3-L1 cells were pre-treated with or without 25 μM tryptamine for 30 min and then treated with 5 nM insulin for the indicated times. The effect of IAA on insulin signaling was evaluated by treating 3T3-L1 cells with or without 100 μM IAA for 60 min followed by 10 nM insulin for the indicated times. To determine the effect of tryptamine on ERK signaling, 3T3-L1 cells were pre-treated with or without ERK inhibitor U0126 (25 μM) for 30 min and then treated with or without tryptamine for 30 min for the indicated times. These cells were further incubated with 10 nM to investigate the role of ERK in the effect of tryptamine on insulin signaling. The effect of tryptamine on the TAAR1-ERK axis was determined by treating 3T3-L1 cells with or without EPPTB (TAAR1 antagonist; 25 μM) for 60 min following by the absence or presence of tryptamine for 30 min for the indicated times. Subsequent incubation with insulin (10 nM; 30 min) was used to establish the TAAR1-ERK axis as a mediator of the tryptamine action on insulin signaling. Tryptamine, U0126 and EPPTB were dissolved in DMSO at 100 mM as a stock solution. For cAMP measurements, 3T3-L1 cells were pre-treated with or without 25 μM of EPPTB for 60 min and then treated tryptamine 25μM for the indicated times.

#### Bacterial strains culture

Tryptamine-producing *Lactobacillus casei* was constructed using the tryptophan decarboxylase (TDC) gene from *Ruminococcus gnavus.* The TDC gene was cloned into the vector and the resulting plasmid was transferred into *L. casei* as previously described (Welker et al., 2015). Successful insertion of TDC gene into the *L. casei* was confirmed by PCR and the production of tryptamine when cultured in an MRS broth containing 0.25% tryptophan. A vector-only strain of *L. casei* was constructed as a control. The engineered *L. casei* TDC+ and vector-only *L. casei* were firstly grown on MRS agar plate containing erythromycin (50μg/mL) and single colonies were isolated and grown in MRS broth at 37 °C overnight. The *L. casei* TDC+ and vector-only *L. casei* were collected from the medium by centrifugation at 3,000 rpm for 10 min at room temperature. *L. casei* inoculums were then prepared in 300 μL of sterilized PBS and then administered to pseudo germ-free mice by oral gavage. Fecal samples were collected daily for measurement of fecal tryptamine level.

### Study methods details

#### Fecal suspension administration

About 10 g of fecal samples were mixed with 5x sterilized 1x PBS (m/v) and homogenized as fecal suspension. HFD-fed mice were orally administered the fecal suspension derived from normal and diabetic monkeys at 4 g/kg daily for 5 days. On day 5, following a 12-h fast, an OGTT was performed to examine the effects of the fecal suspension from monkeys on glucose tolerance.

#### Metabolomics study

About 100 μL of serum and urine were defrosted and extracted with 400 μL of methanol. The samples were vigorously vortexed and centrifuged at 14, 000 rpm at 4 °C for 15 min. About 200 μL of supernatant was transferred to new tubes for LC-MS analysis. For fecal samples, about 150 mg of feces were extracted with 20x 70% methanol (m/v) and then homogenized with steel beads. The samples were then centrifuged at 14, 000 rpm at 4 °C for 15 min. About 200 μL of the supernatant was transferred to new tubes for LC-MS analysis. A pooled quality control (QC) sample was prepared by mixing equal amounts of each sample.

An Agilent ultra-performance liquid chromatography (UPLC) system coupled to a tandem quadrupole-time-of-flight (Q-TOF) equipped with an AJS electrospray interface (G6540A) mass spectrometry was used for untargeted metabolomics. A Waters BEH 2.1×50 mm C18 1.7 μm column with a pre-column was used. The mobile phase used in LC-MS-Q-TOF was A: water with 0.1% formic acid and B: acetonitrile with 0.1% formic acid. The gradients were set as 2-5% B (0-1.5 min), 5-35% B (1.5-6 min), 35-75% B (6-10 min), 75-100% B (10-10.50 min), 100%B (10.50-12 min) and 100%-2%B (12-14 min). The raw MS data were collected and processed using XCMS R package. The data matrix was processed by MetaboAnalyst for multi-variate statistical analysis including PCA/PLS-DA and volcano plot analysis (Chong et al., 2019). The compound identification was performed using online libraries including METLIN (http://metlin.scripps.edu) and HMDB (http://www.hmdb.ca/) based on the MS and MS/MS data.

An Agilent UPLC system coupled to a triple quadrupole (QQQ) 6460 mass spectrometry was used for targeted metabolomics. A Waters BEH 2.1×100 mm C18 1.7 μm column with a pre-column was used. The mobile phase used in LC-MS-QQQ was A: water with 0.1% formic acid and B: acetonitrile with 0.1% formic acid. The gradients were set as 2% B (0-0.5 min), 2-30% B (0.5-4 min), 30-100% B (4-6 min), 100% B (6-8 min), 100-2% B (8-8.1 min) and maintained in 2% B (8.1-10 min). The MS data were collected and processed by the in-house software provided by Agilent. The standards list, MRM transition and retention time were provided in (Supplement Table.S8).

#### Batch culture of fecal samples

About 50 mg of fecal samples were mixed with 20x sterilized 1x PBS (m/v) and homogenized with steel beads. Fecal suspension (20 μL) was inoculated in 2 mL Tryptic Soy Broth (TSB) supplemented with 0.25% tryptophan and incubated overnight under anaerobic conditions at 37°C. After incubation, 100 μL of the medium was then used for quantification of tryptamine by LC-MS analysis.

#### Glucose and insulin tolerance test

The oral glucose tolerance test (OGTT) was used to study the effect of tryptamine on glucose tolerance. Mice were fasted for 12 hours (overnight) and then tryptamine or IAA was orally administered to mice at indicated dosages. After 30 min, mice were orally given glucose at a dosage of 2 g/kg. Blood samples were collected from the tail vein for glucose measurement using Accu-Chek glucose meters at 0, 15, 30, 60, 90 and 120 min after the glucose challenge. To determine the role of ERK and TAAR1 in mediating the effect of tryptamine on glucose tolerance, mice were fasted 12 hours (overnight) followed by intraperitoneal injection of ERK inhibitor U0126 or TAAR1 antagonist EPPTB (1% DMSO in PBS) at the indicated dosages. After 30 min, tryptamine (5 mg/kg) was administered to mice. The mice were then orally given glucose at a dosage of 2 g/kg for OGTT.

For the insulin tolerance test (ITT), mice were fasted for 4 hours before tryptamine was orally administered to mice at indicated dosages. After 30 min, insulin (1 U/kg) was injected intraperitoneally into the mice. Blood glucose levels were measured as per OGTT. For the measurement of serum insulin level, serum was collected 15 min after the mice were treated with tryptamine. To determine the role of ERK and TAAR1 in insulin tolerance, mice were fasted for 12 hours (overnight) and then received ERK inhibitor U0126 or TAAR1 antagonist EPPTB (1% DMSO in PBS) intraperitoneally at indicated dosages. After 30 min, tryptamine (5 mg/kg) was administered to the mice followed by intraperitoneal injection of insulin (1 U/kg) for ITT.

#### Tissue distribution of tryptamine

To determine whether tryptamine can enter the circulation and insulin-sensitive tissues, tryptamine at a dosage of 5mg/kg (dissolved in 0.5% CMC-Na) was orally administered to mice. After 15 min, mice were euthanized by isoflurane and sacrificed by cervical dislocation. Serum, liver, skeletal muscle and white adipose tissues (WAT) were collected and stored at −80°C until analysis.

#### Phospho-proteomics study

Mice were fasted 4 hours and orally administered with tryptamine or vehicle. The WAT tissues were collected and for phospho-proteomics study, whilst other WAT, liver and skeletal muscle tissues were collected and used for the western blotting study. Mice tissue samples used for phosphor-proteomic study were prepared as previously reported (Zhu et al., 2014). The labeled peptides mixtures from each sample was labeled using TMT reagent for MS analysis. An Easy nLC systems (Thermo Fisher Scientific) with an Acclaim PepMap RSLC column (50μm x 15cm) was used to separate the TMT-labeled peptides. The mobile phase used in LC-MS-Orbitrap was A: water with 0.1% formic acid and B: 80% acetonitrile, 20% water with 0.1% formic acid. The elution gradient was set as 0-5 min in 0-6% buffer B, 5-45 min in 6-28% buffer B, 45-50 min in 28%-38% buffer B, 50-55 min 38-100% buffer B and maintained during 50-60 min in 100% buffer B. LC-MS/MS analysis was performed on a Q Exactive Plus Orbitrap LC-MS/MS System (Thermo Fisher Scientific) that was coupled to Easy nLC for 60 min. The mass spectrometer was performed in positive ion mode. MS data was collected according to a data-dependent method to select the most abundant precursor from the survey scan (350-1800 m/z). The normalized collision energy was 30 eV.

The obtained MS/MS spectra were processed by Proteome Discoverer (Thermo Fisher Scientific) and searched using MASCOT engine 2.6. All protein sequences were aligned to the *Mus musculus* database downloaded from UniProt (http://www.uniprot.org). The proteins with a fold change >1.2 or <0.8 and a *p*-value <0.05 were considered as differentially expressed proteins. KEGG pathway annotation was performed using KOALA (KEGG Orthology And Links Annotation) software to identify the significantly enriched pathways by comparing the number of differentially expressed proteins and by matching total proteins to pathways.

#### Protein analysis

Frozen tissues and harvested cells were lysed in RIPA buffer with a protease inhibitor cocktail. For western blotting, the cell lysates and tissue lysates were centrifuged at 15,000 rpm for 15 min at 4 °C. The supernatant was mixed with 5x loading buffer and heated at 98 °C on a dry bath for 10 min. The target proteins were then detected in the samples as per manufacturer instructions. The blots were incubated with HRP-linked anti-rabbit IgG or anti-mouse IgG and reacted with enhanced chemiluminescence.

#### Characterization of tryptamine-producing bacteria

The two reference tryptophan decarboxylase sequences of *Ruminococcus gnavus* (strain ATCC 29149 / VPI C7-9) and *Clostridium sporogenes* (strain ATCC 15579) were downloaded from ENA (A7B1V0 and J7SZ64) (Williams et al., 2014). The identity between these two reference sequences was 26% based on BLASTP. To identify potential tryptophan decarboxylase sequences in the GUT2D dataset, BLASTP was used to align the two reference sequences against the nonredundant microbiome gene catalogue constructed in GUT2D study (Zhao et al., 2018). The alignments were filtered with E-value < 1e-5 and identity > 30%. Repeated measures correlation coefficients between fecal metabolites and abundances of the 5 CAGs were calculated as previous described (Bland and Altman, 1995).

### Statistical analysis

Data were expressed as average and SD or SEM values of at least triplicates. *P*-values were calculated using GraphPad Prism 8 and *p*-values less than 0.05 were considered as statistically significant. Wilcoxon rank-sum test (one tailed or two tailed test) was used to determine the differences of metabolomics data between subjects with and without T2D. Unpaired student’s t-tests or one-way ANOVA were used in other experiments as indicated.

## Supplemental information

**Supplement Figure.S1.**
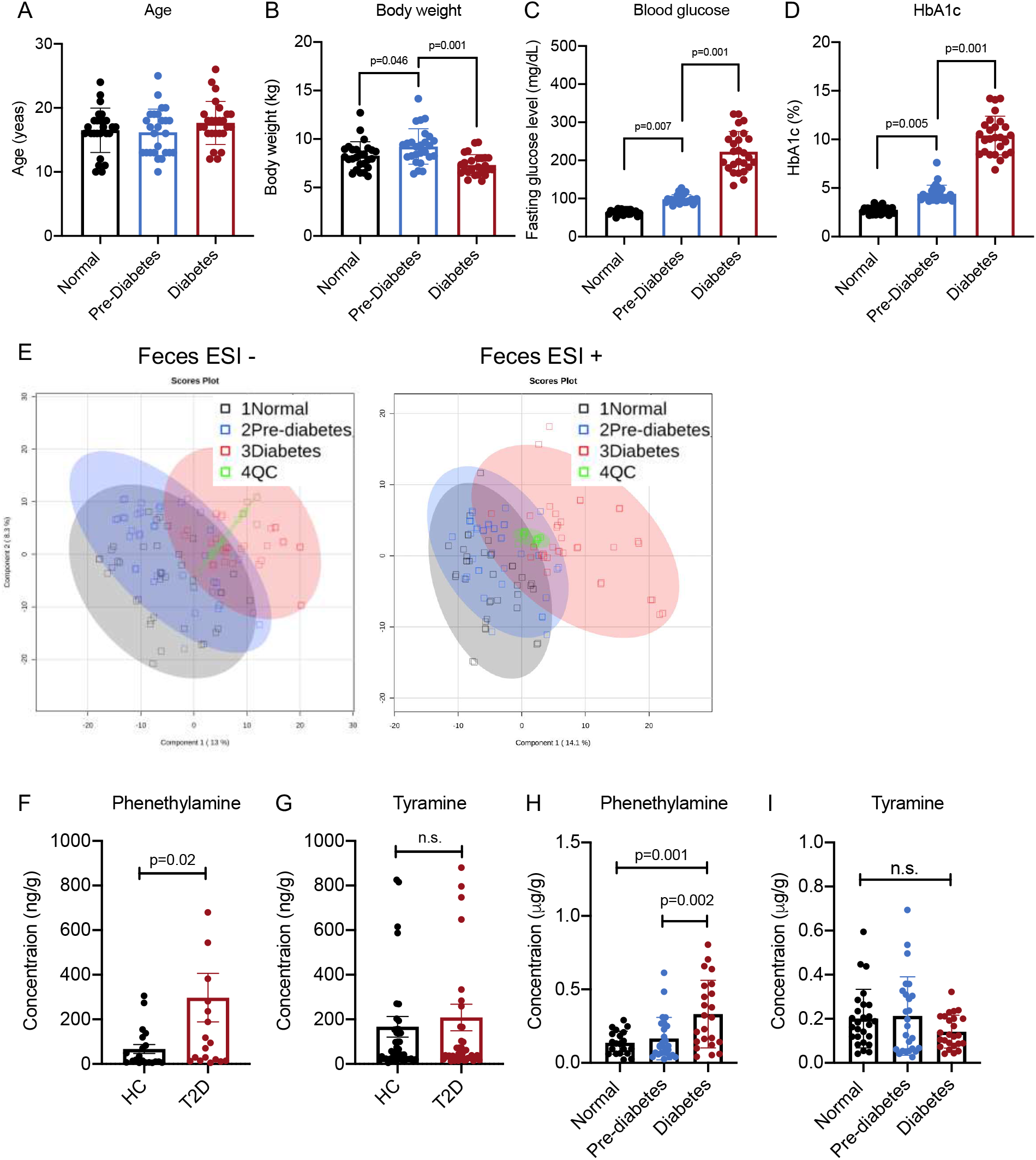
Tryptamine is increased in monkeys with spontaneous diabetes and human with type 2 diabetes. (A-D) Age, body weight, blood glucose and HbA1c index in monkeys with or without diabetes (n=26/per group). (E) Score plot of fecal metabolome in monkeys with or without diabetes with negative and positive (n=26/group). (F-G) Phenethylamine and tyramine levels in fecal samples of human with or without type 2 diabetes (n=25 healthy controls, n=15 diabetic patients). (H-I) Phenethylamine and tyramine levels in fecal samples of age-matched monkeys with or without pre-diabetes and diabetes (n=26/group). Data were presented as mean ± S.D. *P* values were determined by ordinary one-way ANOVA and Student’s t-test.

**Supplement Figure.S2.**
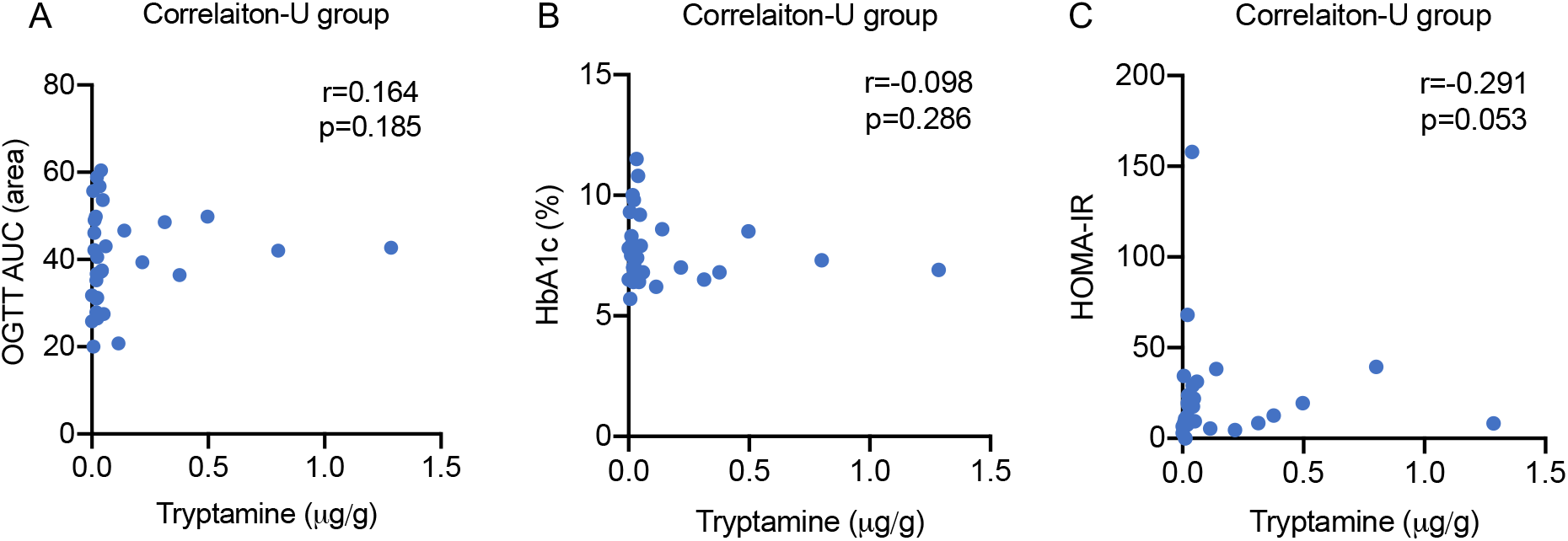
(A-C) Correlation analysis between fecal tryptamine level and OGTT, HbA1c and HOMA-IR indexes in T2D subjects treated with control group (U group; n=16). *P*-values were determined by spearman’s rank correlation.

**Supplement Figure.S3.**
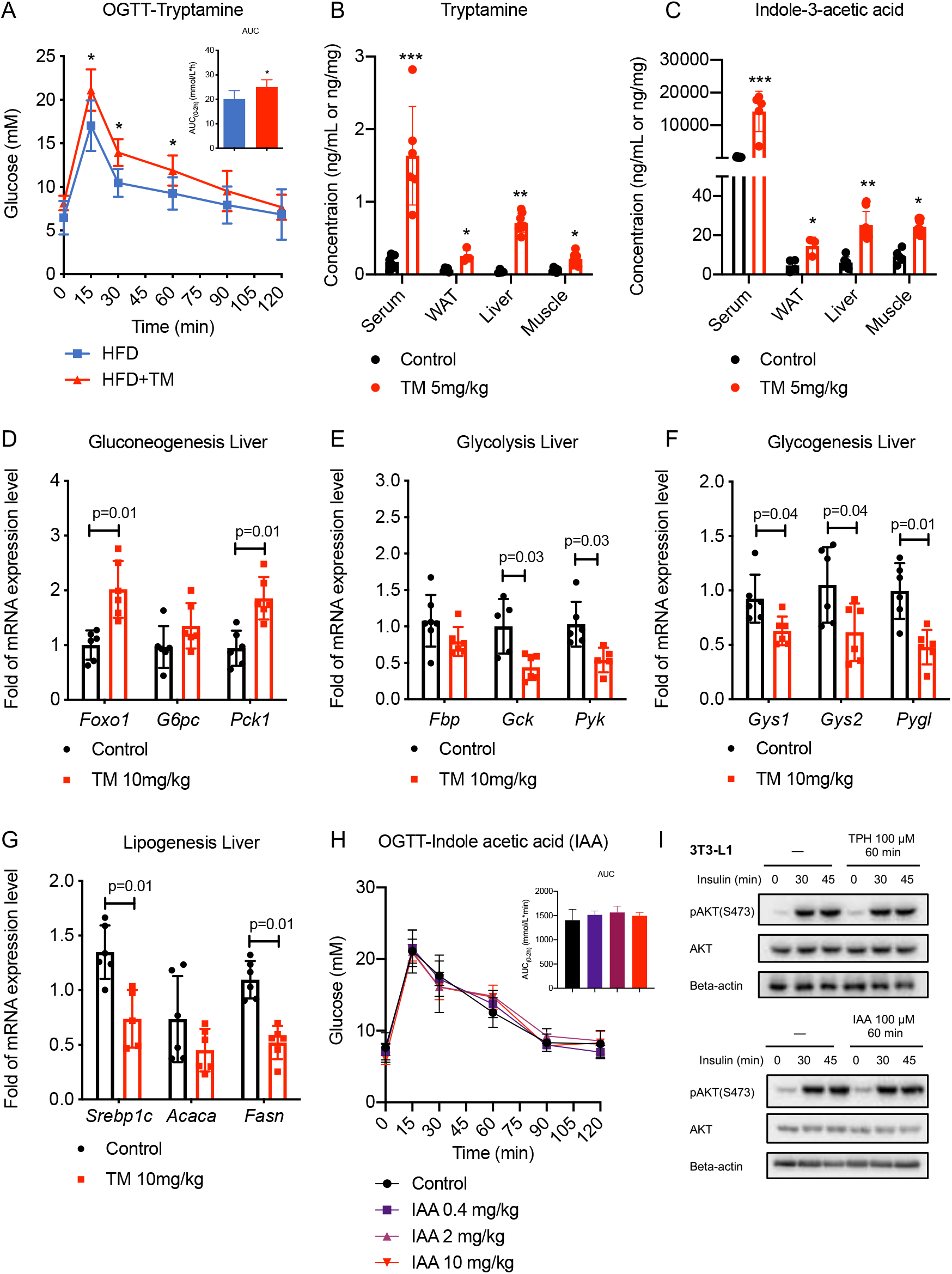
Tryptamine impairs glucose tolerance and insulin sensitivity. (A) Oral glucose tolerance test (OGTT) in high fat diet (HFD) mice after treatment of tryptamine (10mg/kg) (n=6/group). (B-C) Tryptamine and indole acetic acid level in serum, white adipose tissue (WAT), liver and skeletal muscle after administration of tryptamine (5mg/kg) at indicated times (n=6). (D) Oral glucose tolerance test (OGTT) in normal mice after treatment of indole acetic acid (IAA) at indicated dosages (n=6/group). (E) Western blot (and semi-quantification) of tryptophan (TPH) treatment (100μM) and IAA (100μM) on insulin signaling stimulated by insulin (10nM) in 3T3-L1 cells (n=3/group). Data were presented as mean ± S.D. *, p < 0.05, **, p < 0.01 and ***, p < 0.001. *P* values were determined by one-way ANOVA or Student’s t-test.

**Supplement Figure.S4.**
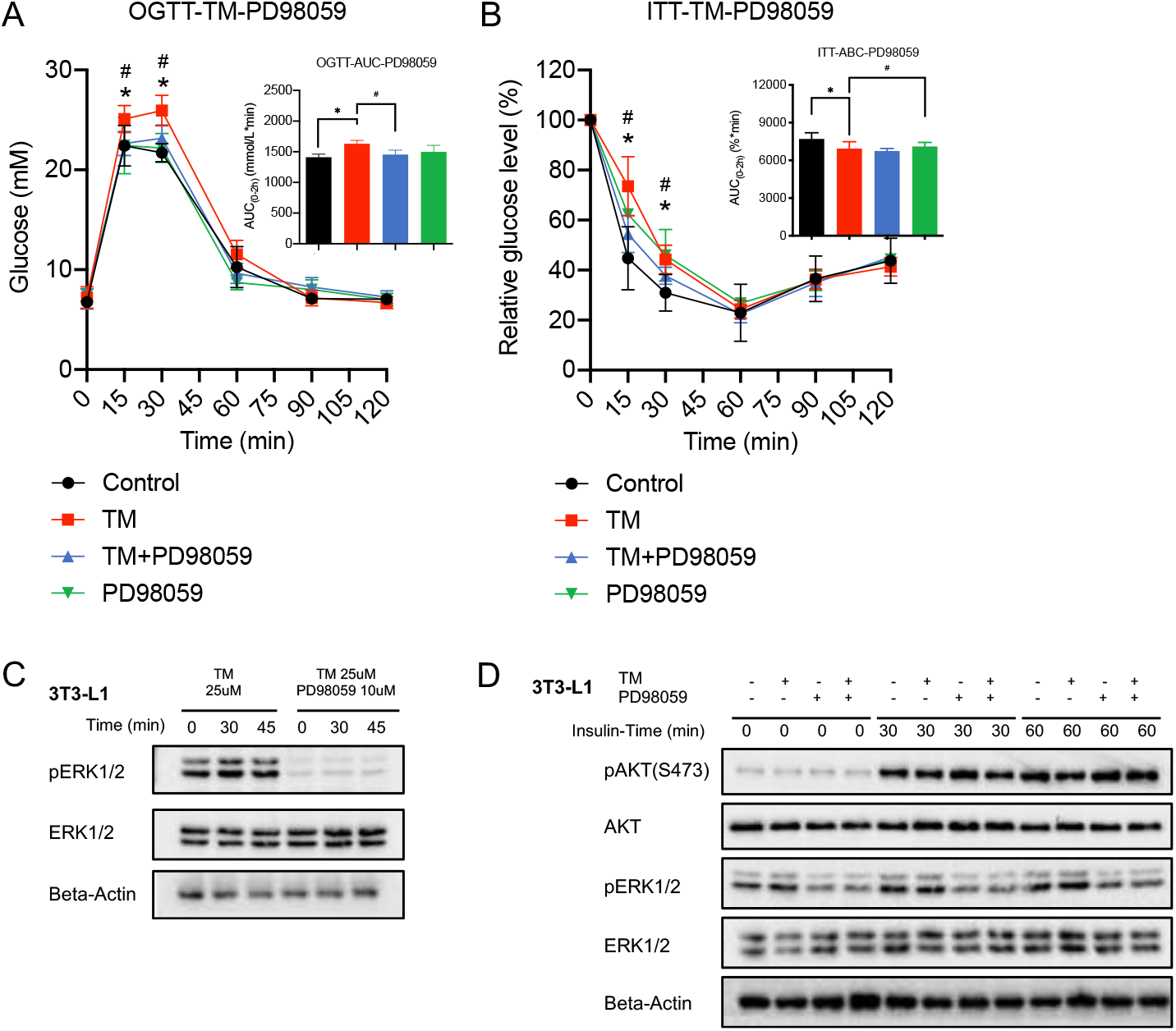
Tryptamine suppresses insulin signaling via ERK activation. (A-B) Oral glucose tolerance test (OGTT) and insulin tolerance test (ITT) in mice after treatment of tryptamine (10mg/kg), ERK inhibitor PD98509 (10mg/kg) or control (1% DMSO in saline) (n=6/group). * comparisons between control group and tryptamine group (10 mg/kg). # comparisons between tryptamine group and tryptamine + ERK inhibitor (PD98059) group. (C-D) Western blot of tryptamine (25μM) and ERK inhibitor PD98509 treatment (10μM) on ERK activation and Akt activation stimulated by insulin (10nM) in 3T3-L1 cells (n=3/group). Data were presented as mean ± S.D. *, p < 0.05, **, p < 0.01 and ***, p < 0.001. p values were determined by one-way ANOVA or Student’s t-test.

**Supplement Figure.S5.**
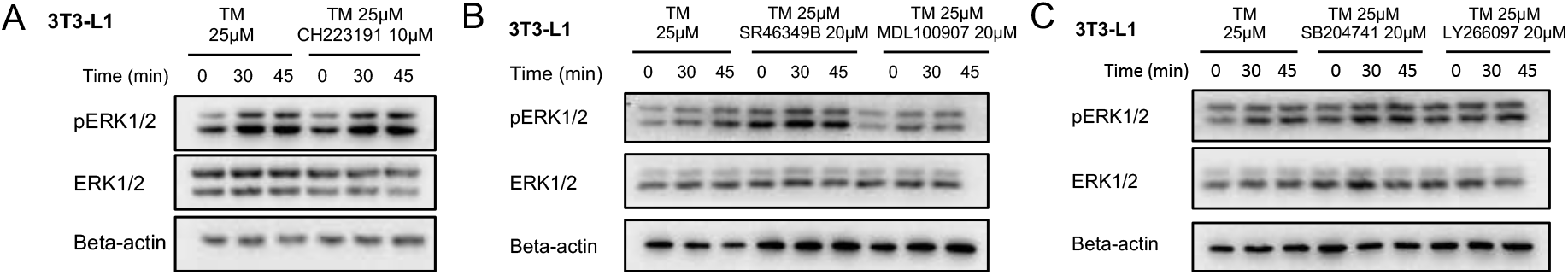
Tryptamine impairs insulin signaling via TAAR1-ERK signaling axis. (A-C) Western blot of tryptamine (25μM), AhR antagonist CH223191 (10μM), 5-HT2A receptor antagonist SR46349B (25μM) and MDL100907 (20μM) as well as 5-HT2B receptor antagonist SB204741 (20μM) and LY266097 (20μM) in 3T3-L1 cells (n=3/group). *P* values were determined by Student’s t-test.

**Supplement Figure.S6.**
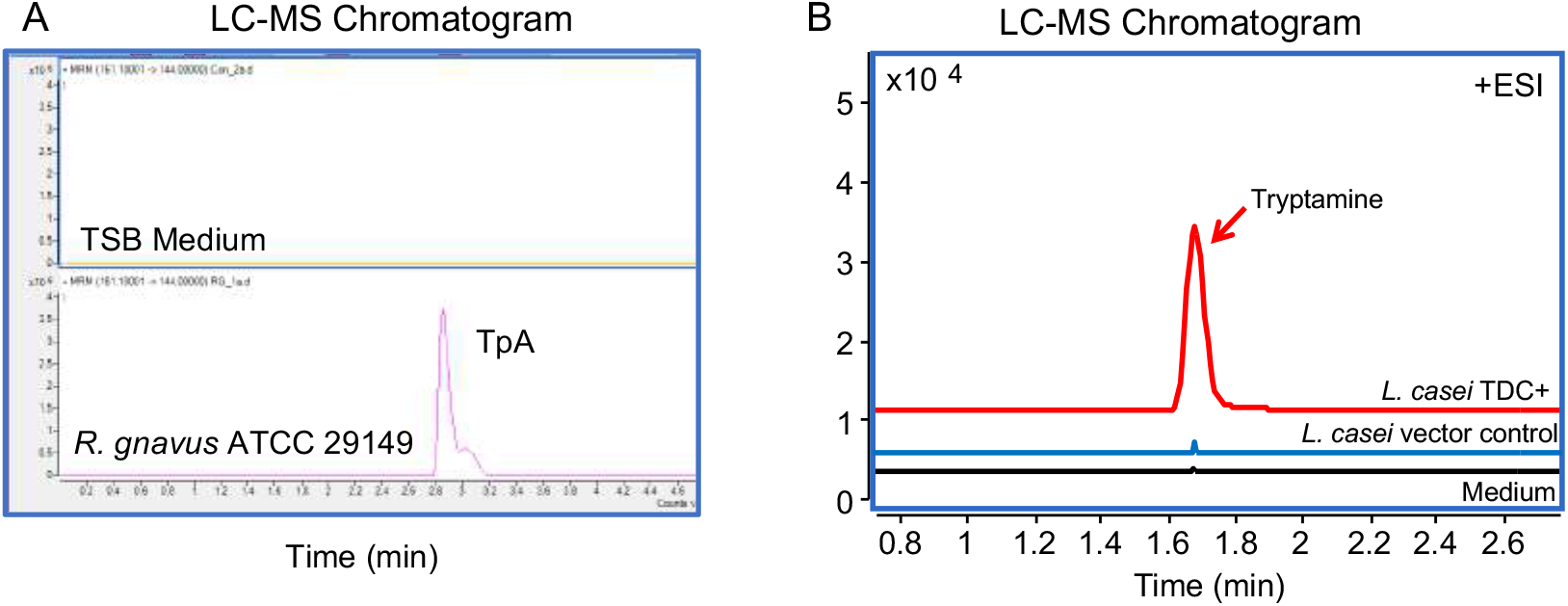
*In vivo* tryptamine production by T2D-associated tryptamine producers and engineered gut microbe impairs insulin signaling. (A) Tryptamine level in culture medium of *R. gnavus* ATCC 29149. (B) Tryptamine level in culture medium of *L. casei* TDC+ and *L. casei* vector control.

**Supplement Figure.S7.**
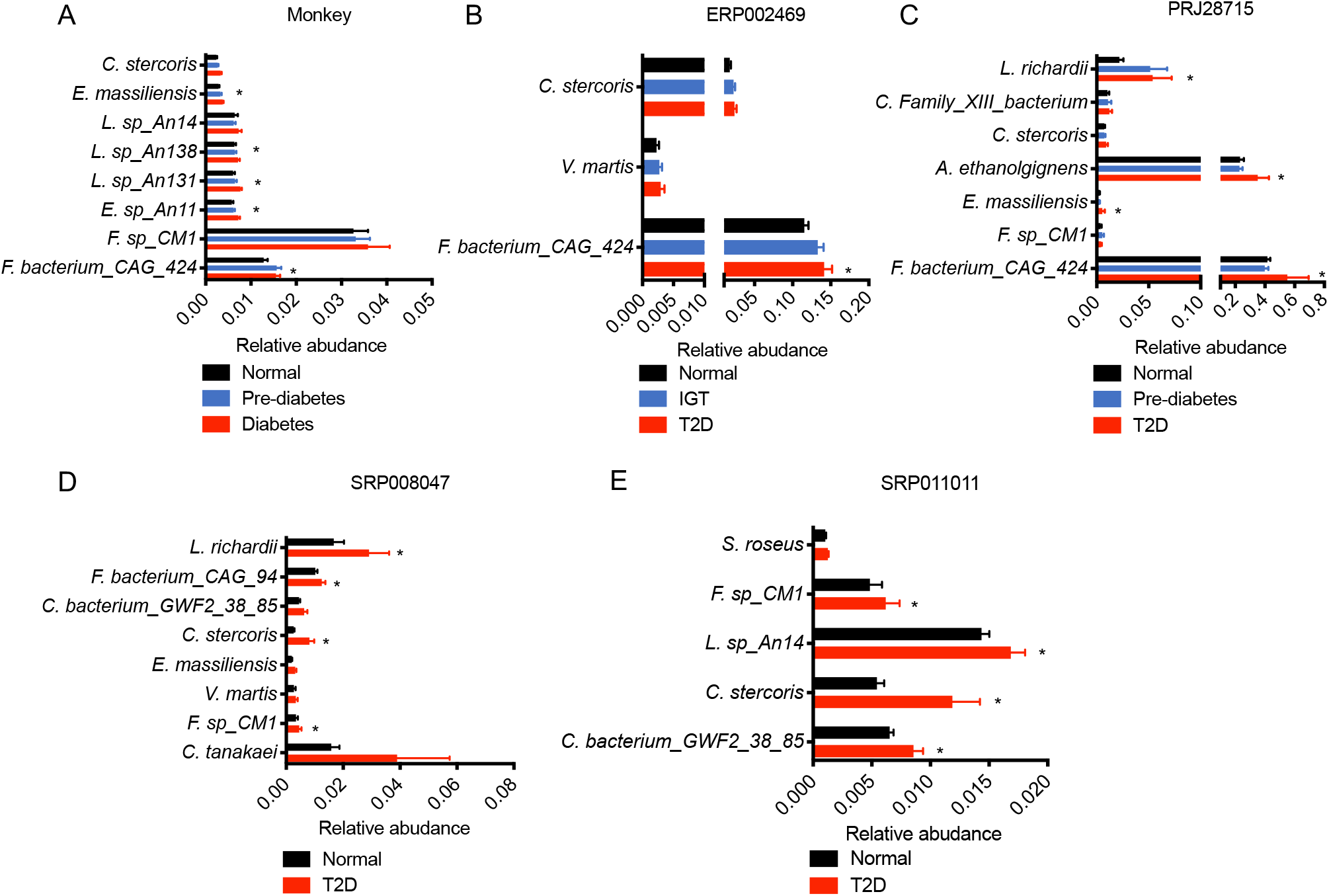
Characterization of T2D-associated tryptamine-producing bacteria. (A-E) The relative abundances of potential tryptamine producing bacteria including *Collinsella stercoris* (*C. stercoris*), *Enorma massiliensis* (*E. massiliensis*), *Lachnoclostridium sp An14* (*L. sp An14*), *Lachnoclostridium sp An138* (*L.* sp An138), *Lachnoclostridium sp An131* (*L. sp An131*), *Eubacterium sp An11* (*E. sp An11*), *Fusobacterium sp CM1* (*F. sp CM1*), *Firmicutes bacterium CAG 424* (*F. bacterium CAG 424*), *Vagococcus martis* (*V. martis*), *Leminorella richardii* (*L. richardii*), *Clostridiales Family XIII bacterium* (*C. Family XIII bacterium*), *Acetivibrio ethanolgignens* (*A. ethanolgignens*), *Firmicutes bacterium CAG 94* (*F. bacterium CAG 94*), *Clostridiales bacterium GWF2 38 85* (*C. bacterium GWF2 38 85*), *Collinsella tanakaei* (*C. tanakaei*) and *Salinicoccus roseus* (*S. roseus*) by references genome blast in monkeys and human subjects with or without T2D. (A) Monkey: Normal (n=20), Pre-diabetes (n=20) and Diabetes (n=22); (B) ERP002469: Normal (n=43), Impaired glucose tolerance (IGT) (n=49); T2D (n=53); (C) PRJ28715: Normal (n=97), Pre-diabetes (n=78) and T2D (n=80); (D) SRP008047: Normal (n=73) and T2D (n=70); (E) SRP011011: Normal (n=105) and T2D (n=109). Data was presented as mean ± S.E.M *, p < 0.05, **, p < 0.01 and ***, p < 0.001. *P* values were determined Wilcoxon one-tailed test.

## Abbreviations

AhR: Aryl hydrocarbon receptor
CAGs: Co-abundance groups
FBG: Fasting blood glucose
FMT: Fecal microbiota transplant
GI: Gastrointestinal
HbA1c: Hemoglobin A1C
HFD: High fat diet
HOMA-IR: Homeostatic Model Assessment for Insulin Resistance
IAA: Indole-3-acetic acid
IR: Insulin resistance
ITT: Insulin tolerance test
IVGTT: Intravenous glucose tolerance test
OGTT: Oral glucose tolerance test
TC: Total cholesterol
TG: Total glycerides
TAAR1: Trace amine-associated receptor 1
Trp: Tryptophan
TDC: Tryptophan decarboxylase
T2D: Type 2 diabetes
WAT: white adipose tissue

## Notes

### Competing Interest Statement

The authors have declared no competing interest.

